# Temperature triggers provide quantitative predictions of multi-species fish spawning peaks

**DOI:** 10.1101/2020.07.10.196923

**Authors:** Emma S. Choi, Erik T. Saberski, Tom Lorimer, Cameron Smith, Unduwap Kandage-don, Ronald S. Burton, George Sugihara

## Abstract

We find a startling correlation (Pearson ρ > 0.97) between a single event in daily sea surface temperatures each spring, and peak fish egg abundance measurements the following summer, in 7 years of approximately weekly fish egg abundance data collected at Scripps Pier in La Jolla California. Even more surprising is that this event-based result persists despite the large and variable number of fish species involved (up to 46), and the large and variable time interval between trigger and response (up to ~3 months). To mitigate against potential over-fitting, we make a true out-of-sample prediction for the peak summer egg abundance that will be observed at Scripps Pier this year.

## Introduction

To comprehend the population dynamics underpinning biodiversity and essential ecosystem services, a heavy emphasis is placed on driving mechanisms, both biotic and abiotic. In marine environments, where fish stocks are of substantial ecological and economic interest, drivers need to be untangled, to inform effective, practical, and sustainable environmental policy. Temperature is a particularly important (and sometimes controversial [1–3]) driver for fish and other marine ectotherm populations. Here we focus on the well-studied relationship between temperature and fish reproduction [4–7]. We find a surprisingly strong correlation between fish-egg- and temperature-dynamics, arising from high-resolution temperature data.

At the seasonal timescale, trends between water temperature and spawning activity have been observed in many fish species [8–17]. Huber and Bengtson [12] found that the gonads of inland silversides (*Menidia beryllina*), a summer spawning species, did not mature to a reproductive level in the absence of increasing water temperatures. In yellow perch (*Perca flavenscens*) both decreasing autumn temperatures and low winter temperatures have been deemed critical in order for gonadal maturation to occur [13, 14]. Additionally, these fish have been manipulated to spawn earlier in the year by increasing the rate of water temperature change [14]. Kayes and Calbert [15] found that for the same yellow perch species, increasing temperature heightened egg production, but even in the absence of a temperature cue endogenous factors could induce spawning. In cyprinid fishes, the initiation of gametogenesis requires low temperatures, but the completion of the process requires increasing temperatures [8]. Other notable studies use degree days, a measure of time based on temperature, to track gonad development from the initiation of vitellogenesis to the onset of spawning [18, 19]. A study done by Henderson et al. [20] demonstrates that the timing of spring transitions and the duration of summer, defined by a temperature threshold, is related to shifts in the center of biomass for multiple species during their seasonal migration to spawning grounds, however, the shifts observed differ by species. These and other varying and apparently complicated specific effects suggest that more general quantitative relationships covering diverse species may be hard to come by.

Despite this, recently a strong quantitative predictive relationship was detected between average winter temperature and average spring-summer egg abundance for a suite of near-shore-spawning species off the coast of southern California [16], which was subsequently supported by out-of-sample data acquired the following year [17]. This relationship is largely explained by colder waters being indicative of largescale upwelling; a process known to supply nutrients to shallower waters [21].

Building on this encouraging result, we re-examine these data of [16] and [17], but now include the additional 2019 data that have since become available, and find a true out-of-sample confirmation of that relationship (Fig. 1C). However, these data, which consist of approximately weekly-sampled, species-identified egg counts of 46 near-shore-spawning species from Scripps Pier since 2013 (see Methods) contain substantial, and possibly important, fine-timescale information that was not considered in the seasonal relationship described in [16]. Though the statistical seasonal association is compelling, it emerges only in large-scale averages. By coupling the full-resolution fish egg abundance time series (Fig. 1A) with daily-averaged sea-surface temperatures (Fig. 1B) from the Southern Californian Coastal Observational Ocean Monitoring System (SCCOOS) dataset, we asked whether finer-timescale temperature dynamics provide information about finer-timescale fish egg abundance dynamics.

**Figure 1.**
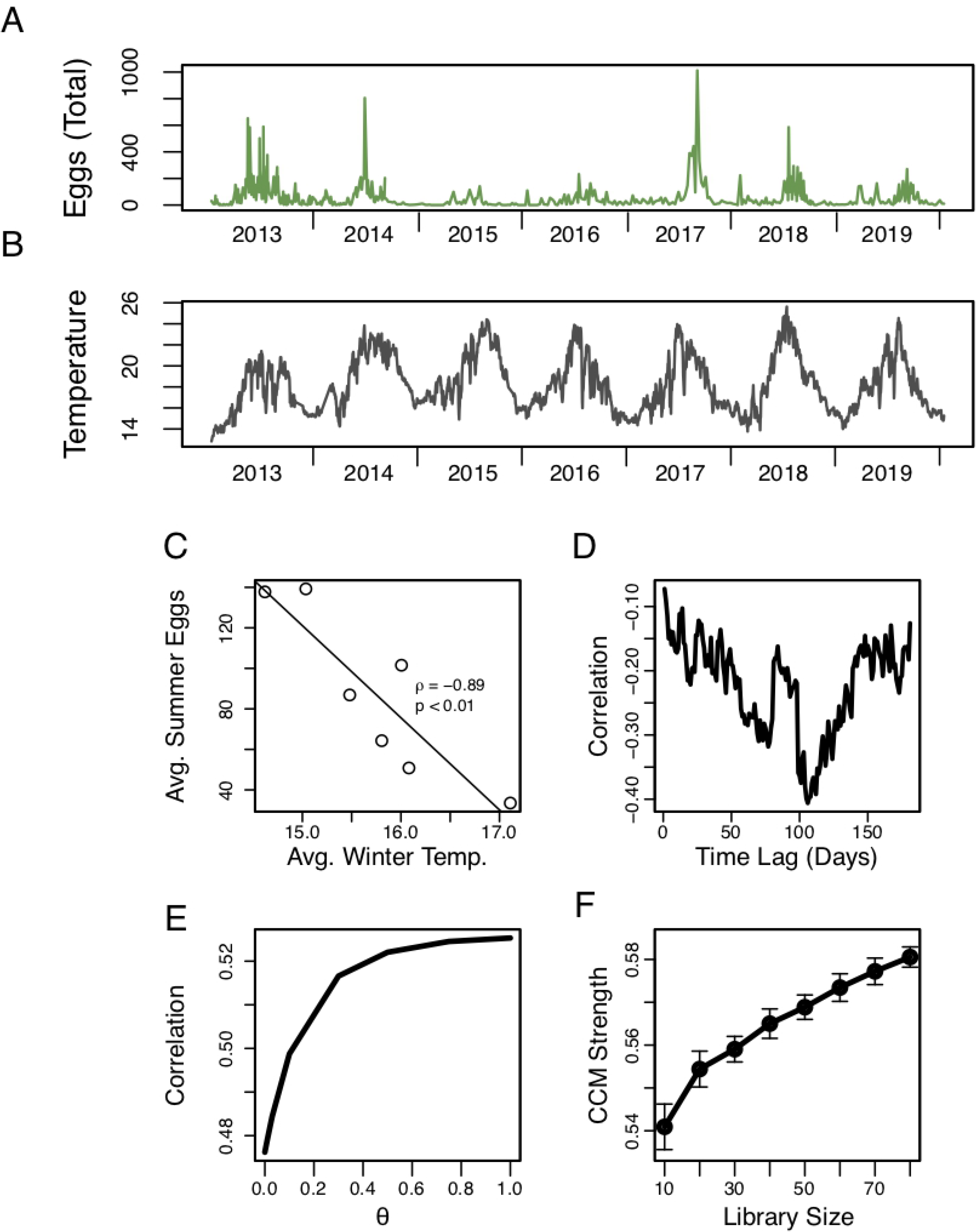
Is there a fine-time-scale relationship between temperature and eggs? A) The total egg abundance in each collection (see Methods) shows substantial variability from year to year in both mean and peak levels. Each fish egg collection was made from the Scripps Institution of Oceanography (SIO) Pier. B) The daily averaged sea surface temperature (SST) in °C at the SIO Pier from data taken every 5-10 minutes from the SCCOOS monitoring station. C) Seasonal averaging reveals the strong negative correlation between the average winter (December – February) SST and the average spring and summer (March – August) egg abundance, identified by [16], with additional points for 2018 and 2019. D) The seasonal correlation breaks down at the daily level; there is no similarly strong correlation between daily winter temperatures and daily egg abundances with time delays ranging from 0 to 180 days. E) The S-Map [22] test for nonlinearity shows that forecasts of egg abundance improve (correlation between predictions and observations) as the nonlinear parameter (θ) is increased, implying that egg abundance shows nonlinear behavior. F) Convergent cross-mapping [23] shows that when using the egg abundance time series to map onto the temperature time series, predictions improve as library size increases, implying there is a dynamic causal effect of temperature on egg abundance.

## Results

The high time-resolution data (Fig. 1 A, B) do not show a linear cross-correlation between the daily spring temperature and lagged daily egg abundance, with only weak relationships across all delays at this fine daily timescale (Fig. 1D). However, in accordance with previous work [24–27], the S-Map test for nonlinearity [22] reveals that the egg abundance is driven by nonlinear processes (forecasts improve as the nonlinear parameter, θ, is increased, Fig. 1E). Further, convergent cross-mapping, a tool for detecting nonlinear coupling in dynamical systems [23] suggests that temperature has a nonlinear effect on egg abundance (converges to ρ = 0.58, n = 295, Fig. 1F). Thus, we expect that a daily timescale relationship may be detectable, just not with linear correlation.

One type of event that stands out in the egg abundance time series (Fig. 1A) is the peak in summer egg abundance. Both the magnitude and timing of the peak egg abundance varies from year to year with no obvious pattern. Previous studies indicate that increasing water temperature may provide a cue for spring and summer spawning species [7, 20, 28, 29]. To ascertain whether a relationship exists between spring temperature increase and peak summer egg abundance, we defined a generic *spring temperature trigger* (STT). Our STT is the maximum of all temperature increases detected within a moving window of length L, as that window moves over the spring season (Fig. 2A, see Methods). This returns a single scalar value for the season, corresponding to a single event with an interpretable characteristic timescale (L). We restricted our analysis of temperature to the spring season (i.e. the season preceding the summer peak) following roughly the causal timescale examined in [16]. By examining a range of possible window lengths, we found a robust relationship around the 1 month timescale, between STT and peak summer egg abundance (L between 3 and 5 weeks, ρ > 0.95, Fig. 2B). This relationship is so remarkably strong (ρ up to 0.98; Fig. 2C) that we feel compelled to share this observation, despite the small number of data points involved (n=7). To mitigate potential overfitting, we offer a prediction for the 2020 peak summer egg abundance that at the time of writing has not yet been measured (Fig. 2C; the largest egg count for the 2020 summer observed at the time of writing is 65 on June 19^th^, 2020). To examine whether this relationship was caused by a general spring warming trend, we repeated the analysis on increasingly smoothed (time averaged) temperature data. We find that the predictive relationship from STT to peak summer egg abundance decreases markedly as temperature becomes increasingly smoothed (Fig. 3), suggesting that there is information about the egg abundance peak specifically at fine time scales.

**Figure 2.**
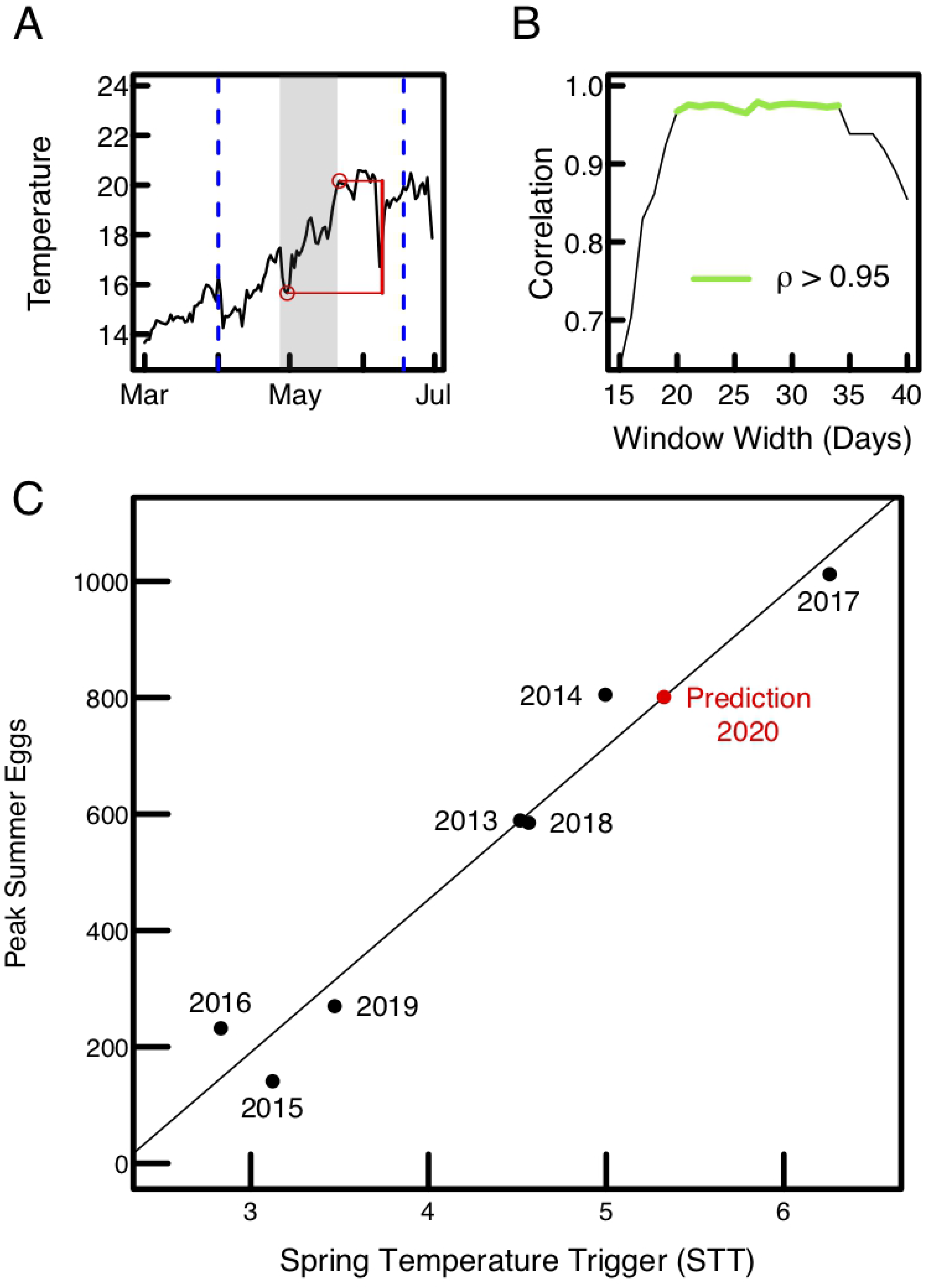
A robust temperature trigger for peak summer egg abundance. A) We define the spring temperature trigger (STT) as the largest temperature increase (denoted in red) detected within a monthly sliding window (gray area) as it moves in daily increments over the spring season (dashed lines; see Methods). B) The relationship between STT and peak summer eggs is robust to the width of the sliding window (widths that produce a ρ > 0.95 are indicated in green). C) The peak correlation between STT and peak summer eggs (June – August) for 2013-2019 (black dots) and predicted value of 801 eggs for 2020 based on the linear regression (red dot).

**Figure 3.**
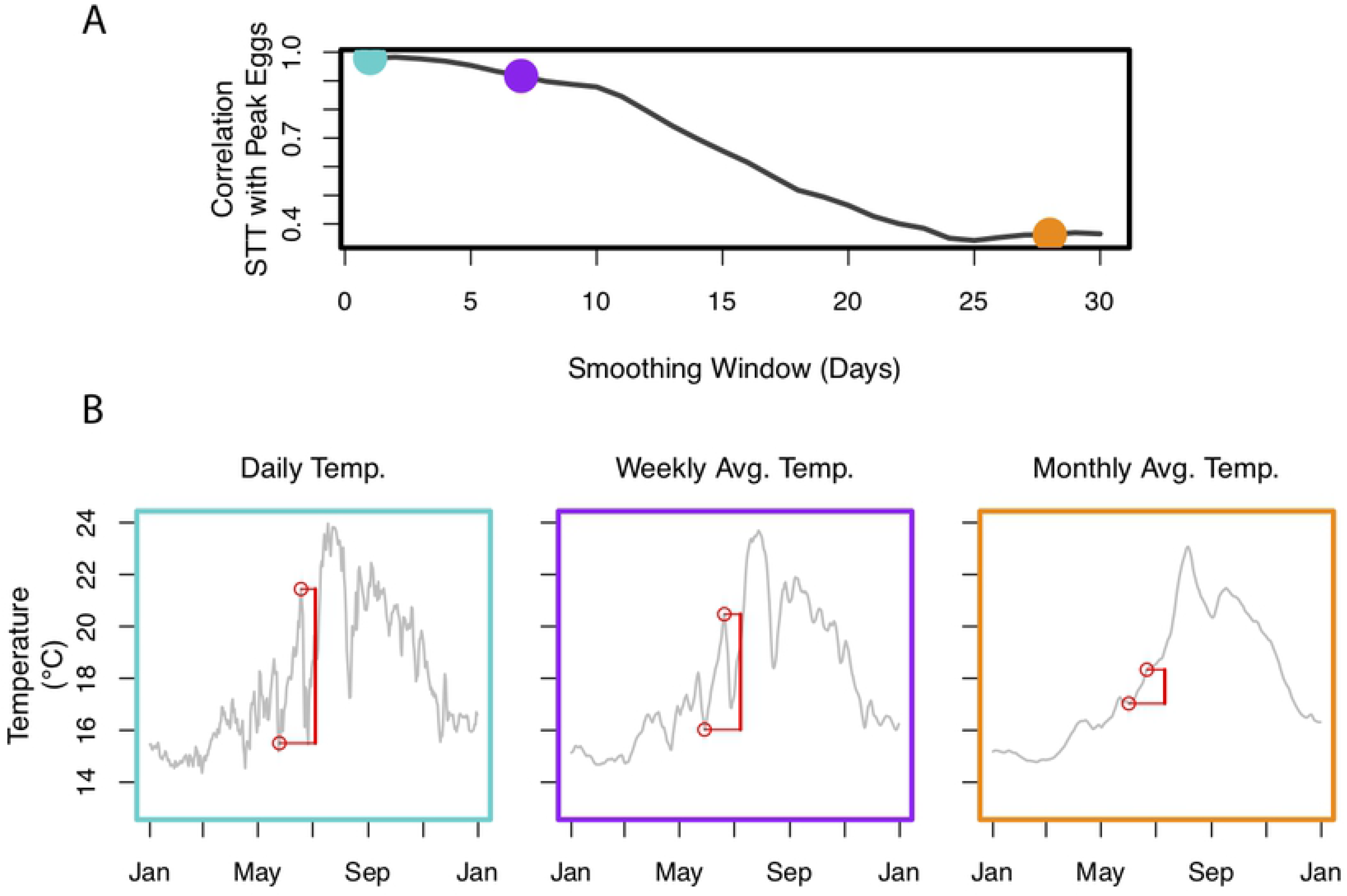
Time averaging obscures the spring temperature trigger. A) The relationship between spring temperature trigger (STT defined over 27 days) and maximum summer fish egg abundance declines as SST is increasingly smoothed across the x-axis (from daily to monthly averages). B) The daily (blue), weekly averaged (purple), and monthly averaged (orange) sea-surface temperature in 2017. Note how the magnitude of the STT (red bars) declines with averaging.

## Discussion

At first glance the positive relationship between STT and peak summer egg abundance appears to contradict the negative relationship found in [16]. However, there are key differences between the two approaches, in both the timing and definition of system input and system response, that may render these apparently different relationships consistent. First, by looking at seasonal averages [16] were focused on broader conditions that may support increased egg abundance driven by, for example, bottom-up processes (upwelling driving increased productivity), while here we consider specific temperature triggers that are blurred in long-term averages (Fig. 3). Second, [16] considers overall winter temperatures and total combined spring-summer egg abundance (as opposed to abrupt changes in spring temperature and peak summer abundance). Despite these differences in the measured quantities, some notable correlations exist amongst them (Fig. S1). Not surprisingly, a significant correlation exists between the peak egg abundance and average egg abundance for a given summer (p =0.02, Fig. S1A). The other correlations shown in Fig S1 could be explained by transitivity; however, it may be possible that winters that are colder on average have more extreme warming events (Figure S1C).

The role of temperature on the timing of the spawning season is of particular interest to living resource managers who utilize forecasts to improve management [30, 31]. Studies have even developed a way to use temperature to track the time from the initiation of gonad development to the onset of spawning by summing the daily mean temperature [18, 19]. The relationship described here does not provide an estimate of the peak spawning timing, but rather only a prediction of the magnitude of the observed peak spawning event.

Physiological temperature triggers for spawning events have been described for multiple species. These studies are often done in laboratory settings (e.g. [15, 32]), or field work focused on sessile invertebrates (e.g. [33–35]). To relate our observations to familiar biological mechanisms, we consider known temperature-related system responses that operate on a time scale similar to what was uncovered here. It has been shown that direct effects of temperature on reproductive processes can be transduced through endocrine pathways [7, 28, 36]. A study conducted in tench (*Tinca tinca*) showed a threshold of 10 degrees C is required to initiate vitellogenesis, but successive ovulation will only occur if the mean daily temperature exceeds 20 degrees C [37]. In salmonids, increasing temperature has been observed to precede increased hormone levels of 17β-estradiol, vitellogenin, and testosterone, all of which are important for successful gametogenesis [38]. Thus, the magnitude of a temperature increase preceding the spawning season may affect a number of endocrine processes leading to the peak abundance of gametes. We conjecture that the presence of thresholds, including on the different endocrine processes underlying gametogenesis, may provide a mechanism by which such an effect as we observe here can be triggered.

In addition to direct individual-level effects, collective effects, such as synchronization of spawning, may also contribute. Maximum daily water temperatures have been shown to have an effect on the synchronization of spawning in damselfishes (*Pomacentridae)* [39]. Sharp temperature increases have been related to shorter and more intense spawning seasons in roach (*Rutilus rutilus*) [29]. Frank and Leggatt [40, 41] documented a relationship between temperature increases, capelin (*Mallotus villosus*) spawning events, and the synchronization of rarer species to those spawning events. However, in our data a preliminary look at the possible influence of synchrony on the magnitude of summer peaks does not appear to support this potential inter-species-level mechanism (Fig. S2). Clearly, synchrony effects will be difficult to definitively separate from individual-level physiological effects (particularly in our in-situ data), and so we leave the problem of distinguishing these two mechanisms as an open question for future work.

## Materials and Methods

### Convergent Cross Mapping

Convergent cross mapping was performed using the block_lnlp() function in rEDM *v*0.7.3. The embedding was made with the raw egg abundance data and the daily-averaged SCCOOS temperature time series.

When performing CCM, the variable being predicted is the one being tested as a causal driver. As such, we used the egg abundance time series to predict temperature (thus measuring temperature’s effect on egg abundance). Although temperature data existed for every day, eggs were collected at inconsistent intervals, typically ranging from one collection every 2-5 days. Thus, in order to make a proper embedding, we filtered both temperature and egg abundance time series to only include temperature values for which a collection occurred on a given day, 6-8 days prior, and 13-15 days prior as well. This gave us a 3-dimensional embedding for fish eggs, with time lags of about 1 week, with accompanying temperature values.

Because both egg abundance and temperature are strongly seasonally driven, we needed to make sure we were not identifying shared information in the two variables driven by seasonality. To account for this, nearest neighbor selection only considered time points that were within 90 calendar days for our target prediction. Without doing this, increased library size will only increase the amount of seasonal information resolved in the embedding rather than actual causal inference.

Libraries of potential neighbors (points within 90 calendar days of the date of the target) were generated at random for each predicted point. Library sizes ranged from 10 - 80 points (increasing by increments of 5). Once the library was randomly generated, the nearest 4 neighbors (E+1, see (22)) in state space were selected and used to make a prediction. After a prediction was made on each temperature value, Pearson’s correlation was calculated between observed and predicted values. This process was repeated 50 times for each library size.

### STT calculation

As described in the text and illustrated in Fig. 2, STT was calculated by allowing the last day of the sliding window to move over the spring interval defined as April 1st to June 18th.

### Sample collection

*V*ertical plankton tows (approximately weekly) were conducted off of the Scripps Pier (32.8328° N, −117.2713° W) from 2013 to 2019. A 1-meter diameter net with 505 micron mesh and a bottle attached to the cod end was lowered to the seafloor, approximately 5 meters, and out of the water 4 times, sampling a total of ~16 cubic meters. The net was then rinsed by lowering it into the water until the top of the net touched the surface and then raised back out. While there is some variation in the volume of water being sampled (e.g., water depth changes with tide, but since eggs are buoyant, most are collected near the surface) and the time of day samples occur, we do not expect a strong influence of either on the peak summer egg abundance. Currents could affect sample volume but are rarely strong in the summer [42] and are therefore less likely to skew the value of peak eggs. Second, the eggs captured at the pier all originated 0-3 days before the collection occurred since in Southern California water temperatures, most fish eggs all hatch within 72 hrs [43] hence any eggs from a spawning event preceding the collection by up to 3 days could still be represented in our sample, depending on precise spawning location and currents. Using real-time current velocities, retrospective modeling found that most eggs collected at the Pier site likely originated within a few kilometers of the collection site [43].The contents of the cod end were concentrated through a 330 micron mesh screen and then sorted under a microscope at 10X. The morphologically distinct eggs of the Northern anchovy (*Engraulis mordax*) and the Pacific sardine (*Sardinops sagax*) were counted and stored separately. The remaining fish eggs were counted and grouped in a 1.5 mL microtube containing 95% ethanol until further processing (identification by DNA barcoding).

## Acknowledgments

This work was supported by DoD-Strategic Environmental Research and Development Program 15 RC-2509 (GS), NSF DEB-1655203 (GS), NSF ABI-1667584 (GS), DOI USDI-NPS P20AC00527 (GS), the Scripps Institution of Oceanography Postdoctoral Fellowship (TL), the McQuown Fund and the McQuown Chair in Natural Sciences, University of California, San Diego (GS). Fish egg collection and identification was supported in part by the Richard Grand Foundation and the California Ocean Protection Council R/OPCSFAQ-12 (RB).

---

**Figure S1**. Correlation between multiple variables studied. The strongest correlation we found was between STT and peak egg abundance (maximum correlation of 0.98). However, as found by [16], a strong, negative correlation exists between average winter temperatures and average summer egg abundance. Not surprisingly, there is also a strong correlation between peak egg abundance and average egg abundance for a given summer (A). Due to transitivity, there is also a strong correlation between average summer eggs and spring temperature triggers (B). Weaker correlations also exist between average winter temperatures and the finer scale temperature triggers (C) and peak summer egg abundance (D), however these are much weaker relationships (p> 0.1).

**Figure S2**. Inconclusive evidence of between-species synchrony in peak summer egg abundance. Shannon diversity (base *e*) appears to be higher for peaks with lower abundance. We note however, that this null result may be influenced by amplification or sequencing failures that do not allow us to identify every egg in the sample to species (a limitation that does not affect total egg counts).

